# VARION: A Network Propagation Framework for Individual Patient Somatic Mutation Interpretation in Cancer Molecular Subtyping

**DOI:** 10.64898/2026.09.11.751075

**Authors:** Taesoo Kwon, Young-Gyu Park, Jong-Gwon Choi

**Author notes:** Correspondence to:* Professor Jong Gwon Choi, Oncology-Hematology, Konyang University Hospital, 158 Gwanjeodong-Ro, Seo-Gu, Daejeon, Republic of Korea.

## Abstract

Accurate molecular subtyping of individual cancer patients from somatic mutation data remains a challenge in precision oncology research. Existing network-based stratification (NBS) methods treat all mutations equivalently, require full-cohort batch processing, and do not demonstrate generalization to independent datasets without retraining. To address this, we present variant interpretation via the adaptive network pRopagatION (VARION), which integrates population-level variant constraint scoring with protein–protein interaction (PPI) network topology. The Adaptive Topology-aware Random Walk with Restart (ATR-RWR) algorithm weights each mutated gene by φg = √(GIS(g) × ρtopo(g)), where GIS (Gene Intolerance Score) reflects population-level functional constraint, propagated across a shared PPI network; subtype assignment then uses cosine similarity to TCGA-derived reference centroids, enabling real-time single-patient classification. Across ten TCGA cancer cohorts (n = 2,417), VARION achieved 77.7% accuracy for ovarian cancer (OV), 69.5% for glioblastoma (GBM), 90.2% for cholangiocarcinoma (CHOL), and 75.4% for gastric cancer (STAD). A controlled benchmark applying two alternative clustering methods (PyNBS; a dense autoencoder) to identical ATR-RWR propagation matrices recovered no significant driver enrichment (OR = 1.79 and 1.52, n.s.), versus OR = 144.29 (p = 1.77×10⁻¹²) for VARION, confirming that the GIS-weighted centroid architecture, not propagation alone, drives performance; generalization without retraining was further confirmed in two independent cohorts (ICGC CCA, n = 396; PCAWG, n = 110; OR = ∞, p < 5×10⁻⁹). Together, these results indicate that VARION’s GIS-weighted centroid architecture enables individual-patient molecular subtyping that outperforms existing NBS and graph-learning clustering approaches, with high sensitivity for clinically actionable rare subtypes and robust cross-platform generalization.

## 1. Introduction

Precision oncology is fundamentally based on the recognition that patients sharing the same anatomical cancer diagnosis may harbor biologically distinct diseases requiring different therapeutic strategies. Histopathological examination has traditionally provided the foundation for cancer classification, using tumor morphology together with immunohistochemistry (IHC) to establish tumor type and lineage. This approach remains clinically indispensable, achieving a median concordance of 86% for ovarian carcinoma, which improves to approximately 90% with IHC [1]. However, histological similarity does not necessarily imply molecular equivalence; high-grade serous ovarian carcinoma (HGSOC) is recognized histologically as a single entity, yet transcriptomic analyses have consistently revealed distinct molecular states, including immunoreactive, differentiated, proliferative, and mesenchymal subtypes, with differences in tumor biology and clinical outcomes among these groups [2–4].

This distinction is becoming increasingly important because molecular stratification is no longer confined to retrospective cancer taxonomy but is progressively influencing therapeutic development and clinical trial design. Biomarker-enrichment strategies are now routinely incorporated into oncology trials to identify populations most likely to benefit from targeted therapies [5]. In the phase III RUBY trial, for example, dostarlimab produced a substantially greater treatment effect in endometrial tumors with deficient mismatch repair or microsatellite instability-high status than in the overall population [6]. In glioma, IDH status has become integral to disease classification following the phase III INDIGO trial, which demonstrated the clinical benefit of vorasidenib specifically in IDH1- or IDH2-mutant grade 2 glioma [7]. Such examples indicate a broader transition in oncology from treatment based predominantly on anatomical or histological diagnosis to therapeutic strategies designed around molecularly defined patient populations.

Cancer molecular subtyping has transformed our understanding of tumor biology and has enabled increasingly stratified treatment strategies [2, 3]. Large-scale multi-omics initiatives such as The Cancer Genome Atlas (TCGA) have been central to this transformation, integrating genomic, transcriptomic, epigenomic, and copy-number information to reveal biological structures that cannot be resolved by morphology alone [2]. However, the same dimensional richness that gives multi-omics its biological power also creates major barriers to routine clinical implementation, including the need for multiple assays and sample types, high cost and computational burden, and challenges in achieving reproducible, clinically interpretable results across institutions [8–10]. An illustrative example is the recent case of a technology executive whose recurrent osteosarcoma was characterized using multimodal genomic, transcriptomic, and single-cell profiling to inform an individualized neoantigen vaccine strategy [11, 12], underscoring both the promise and the practical burden of comprehensive multi-omics characterization for routine care.

Consequently, a major unmet need is not simply to generate more molecular information, but also to extract clinically meaningful patient states from molecular data that are already routinely available. Population-level molecular classifications, such as those generated by TCGA, were primarily developed to discover biological structures within large cohorts and do not directly provide a practical framework for assigning a newly presented individual patient to a molecular subtype in real time. Somatic mutation profiles represent an attractive starting point for overcoming this gap, because they are already generated by standard clinical NGS panels, are comparatively scalable, and do not require additional RNA extraction or transcriptomic profiling. A computational framework capable of transforming these sparse mutation profiles into biologically informative molecular states could therefore extend the benefits of molecular subtyping to individual patients, while substantially reducing the experimental complexity of multi-omics profiling.

Network-based stratification (NBS) [13] addresses the sparsity of somatic mutation data by propagating mutation signals through protein–protein interaction (PPI) networks. Subsequent refinements, including PyNBS [14], struc2vec-based approaches [15], and graph neural-network methods such as DeepGraphMut [16], have improved subtype stability and survival prediction. Despite these advances, three fundamental limitations persist in the existing mutation-based stratification methods.

First, the existing methods generally encode somatic mutations without explicitly weighing their biological importance at the variant level. DeepGraphMut [16], a recent graph-learning approach for mutation-based cancer stratification, represents mutation status as a binary input, thereby not explicitly incorporating variant class or population-level evolutionary constraints into the initial mutation signal. Second, these approaches are predominantly cohort-oriented algorithms in which samples are analyzed jointly, limiting their direct use for the real-time classification of a single newly presenting patient. Third, validation has largely relied on TCGA-derived datasets and the ability to assign independent patients from external cohorts to predefined molecular states without model retraining remains insufficiently demonstrated.

VARION was developed to address these limitations simultaneously. First, the Gene Intolerance Score (GIS) incorporates a population-level variant constraint as a signal-quality weight before network propagation, enabling biologically differentiated weighting of somatic alterations. Second, centroid-based assignment against precomputed TCGA reference centroids enables the deterministic classification of an individual patient without cohort reprocessing or model retraining. Third, we evaluated the generalization in independent ICGC [17] (n = 396, 10 countries) and PCAWG [18] whole-genome sequencing (n = 110) cohorts without retraining. We additionally performed a controlled benchmark designed to separate the contribution of centroid-based classification from that of network propagation by comparing VARION with PyNBS and RWR- AE, using identical ATR-RWR propagation matrices. Whereas existing knowledge bases and molecular diagnostic frameworks primarily annotate individual variants, retrieve therapeutic evidence, or rely on RNA-based or multi-omics classifiers, VARION is designed to perform individual-patient molecular subtype assignment directly from somatic mutation calls, without requiring RNA expression profiling.

## 2. Methods

### 2.1 Data sources

Somatic mutation data (MAF format) for ten TCGA projects were obtained from the GDC Data Portal. The shared PPI network comprised 2,291 genes and 204,453 edges (STRING v11.5 [19] combined score ≥ 400, BioGRID v4.4 [20]). Independent validation cohorts: ICGC CCA 2017 (chol_icgc_2017; n = 396) and PCAWG OV (Ovarian serous cystadenocarcinoma) (pancan_pcawg_2020; n = 110), both retrieved from cBioPortal [21, 22].

### 2.2 Gene-level weight (φ_g_) and ATR-RWR propagation

The VARION algorithm proceeds through three stages (Figure 1): (Stage 1) gene-level weighting, (Stage 2) network propagation via the Adaptive Topology-aware Random Walk with Restart (ATR-RWR), and (Stage 3) centroid-based subtype assignment.

**Figure 1.**
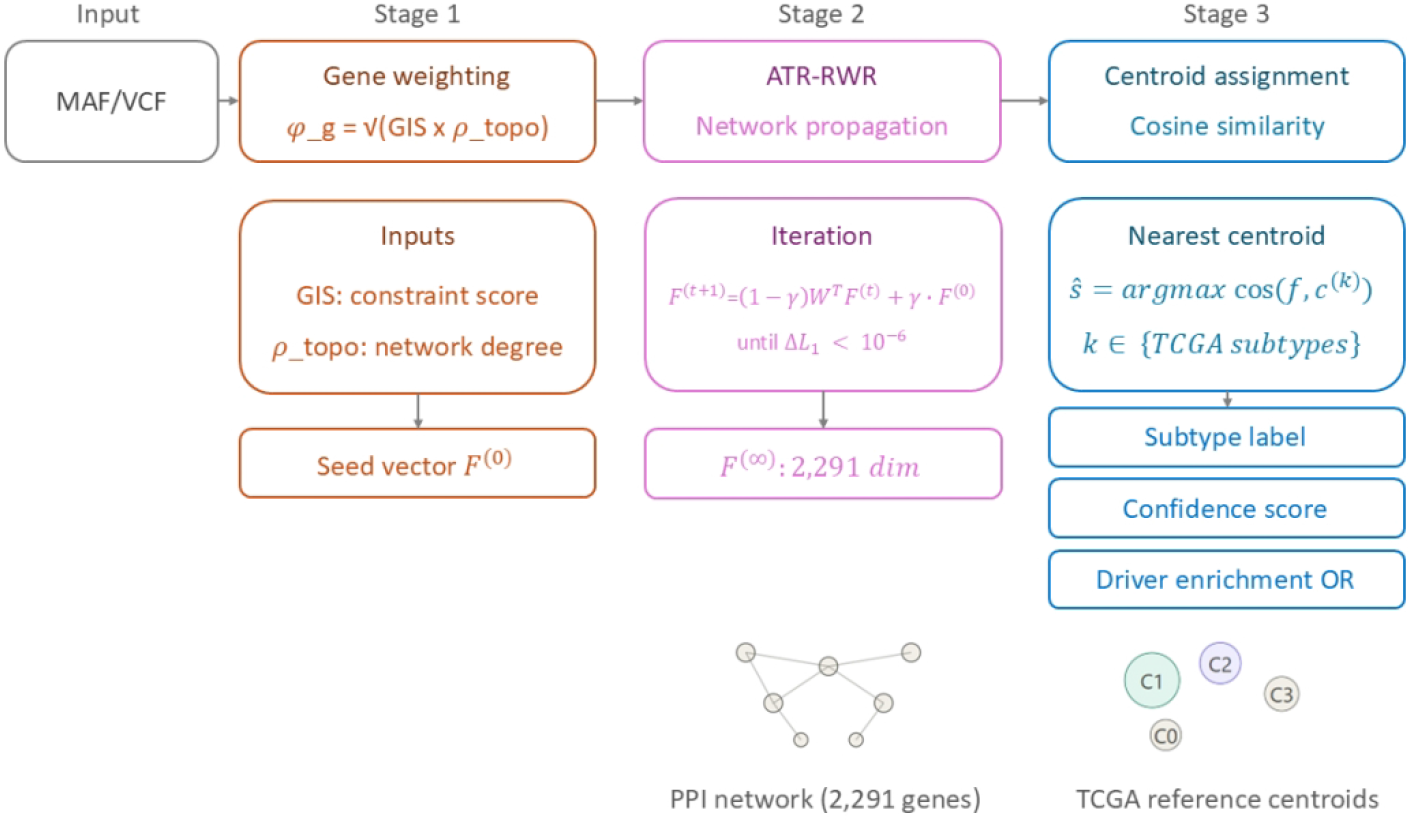
Overview of the VARION computational pipeline. Somatic mutations are weighted by φ_g_ = √(GIS(g) × ρtopo(g)) and propagated through the shared PPI network via ATR-RWR. The resulting propagation vector is assigned to the nearest TCGA reference centroid via cosine similarity, yielding a subtype label, confidence score, and driver gene enrichment report.

#### Stage 1 : Gene-level weighting

For each gene *g* in the shared protein–protein interaction (PPI) network (*N* = 2,291 genes), a composite weight *φ_g_* is defined as the geometric mean of two independently computed gene-level scores:

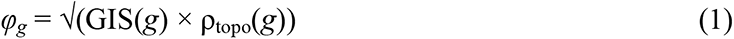

GIS(g) ∈ [0, 1] is a proprietary, patent-protected gene intolerance score derived from population-level variant-intolerance metrics (see Data availability); higher values indicate stronger evolutionary constraints and greater likelihood of functional consequences.

ρ_topo_(*g*) is the network-topology term, computed as the equal-weight mean of two min-max-normalized centrality measures on the shared PPI graph:

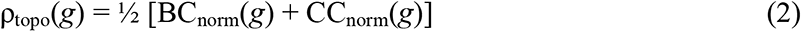

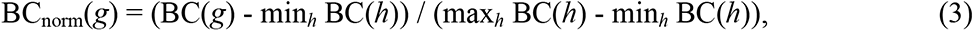

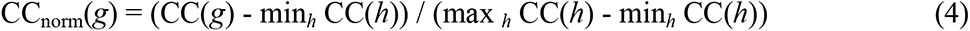

BC(g) and CC(g) denote betweenness and closeness centralities, computed once on the network topology independent of patient data; minh and maxh denote the minimum and maximum over all genes h. ρtopo(g) ∈ [0, 1] captures the overall network centrality without favoring either measure.

The geometric meaning in Equation (1) keeps φg low whenever either term is low, penalizing genes that are network-central but population-tolerant, or vice versa, while bounding φg within [0, 1]. For a given patient, let *M* denote the set of genes carrying a somatic mutation. The initial seed vector *X* ∈ ℝ^2291^ is constructed elementwise, asfollows:

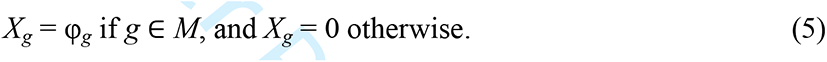

and is then L1-normalized,

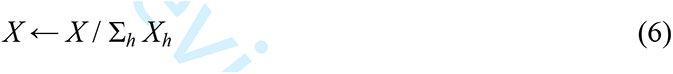

such that X is a probability distribution over the 2,291 network genes, weighted by *φ_g_*. *X* serves as both the initial condition *F*^(0)^ and fixed restart distribution used in Stage 2.

#### Stage 2 : ATR-RWR network propagation

Stage 2 propagated the seed vector *X* across the shared PPI network using an Adaptive Topology-aware Random Walk with Restart (ATR-RWR). The propagation vector was iteratively updated asfollows:

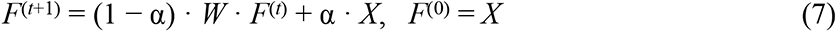

*F*^(*t*)^ ∈ ℝ^2291^ is the propagation vector at iteration *t*, and *W* is the degree-normalized (column-normalized) adjacency matrix of the PPI network, defined element-wise as

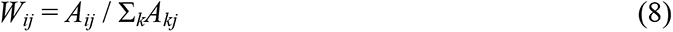

where *A* is a binary PPI adjacency matrix. Column-normalization ensures that *W* · *F^(t)^* redistributes mass evenly across neighbors, that is, *W* is a standard column-stochastic transition operator.

The scalar α = 0.85 is the restart probability (conventional range 0.7-0.9): with probability α, the walk resets to *X*, and with probability (1 - α), it diffuses one step via *W*, keeping the signal concentrated near the seed genes and their close network neighbors.

The iteration of Equation (7) continues until the propagation vector converges, which is defined as the L1 distance between successive iterates falling below a fixed tolerance:

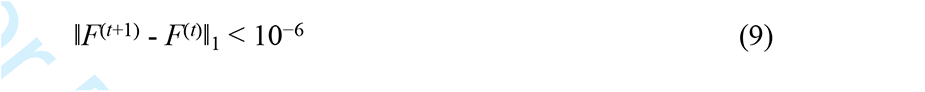

Convergence under this criterion is typically reached in fewer than 100 iterations, with a hard cap of 1,000 iterations imposed to guarantee termination. The converged vector, denoted *F*^(∞)^, is a 2,291-dimensional real-valued vector summarizing the propagated influence of the patient’s mutation profile across the entire shared network and constitutes the patient-level feature representation used for subtype assignment in Stage 3.

When Stages 1-2 are applied to an independent cohort whose original variant calls cover a different, typically smaller set of genes, the resulting propagation vector is reindexed to the full shared gene space before Stage 3, with any absent gene set to zero. This guarantees that *F*(∞) always occupies the same 2,291-dimensional coordinate system regardless of the originating cohort or sequencing platform, allowing centroid comparison in Stage 3 to be performed consistently across cohorts.

#### Stage 3 : Centroid assignment

Stage 3 assigns each patient to a molecular subtype by comparing their propagation vector *F*^(∞)^ against a set of precomputed reference centroids using cosine similarity.

For a cancer type with subtypes k ∈ {1, …, K} (K typically 3-4), the reference centroid for subtype k is the mean propagation vector of the training set patients assigned to that subtype:

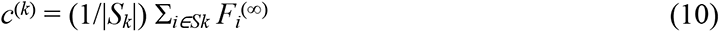

where Sk is the training patient index set for subtype k. During cross-validation, centroids are recomputed independently within each training fold and applied to the corresponding held-out fold; therefore, no held-out patient’s data contributes to its own comparison centroid.

Each query patient is assigned a subtype whose centroid maximizes cosine similarity to its own propagation vector:

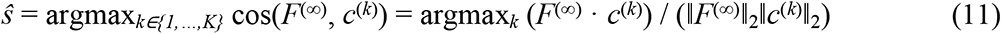

Cosine similarity was preferred over Euclidean distance because it is independent of vector magnitude, which reflects the tumor mutational burden rather than subtype identity. In addition to the subtype label *ŝ*, VARION reports a confidence score, taken as the cosine similarity value cos(*F*^(∞)^, *c*^(*ŝ*)^) attained by the assigned subtype, and a post-hoc driver-gene enrichment odds ratio, computed by Fisher’s exact test, comparing the frequency of a candidate driver mutation among patients assigned to a given subtype against its frequency among all other patients in the same cancer cohort (Section 2.4).

### 2.3 Benchmark comparison

To isolate VARION’s centroid architecture of VARION from the propagation step, two alternative clustering methods were applied to identical ATR-RWR propagation matrices for OV and CHOL (Cholangiocarcinoma): (1) NMF+K-means (PyNBS [14] ; k = number of subtypes, 2,000 iterations, nndsvda initialization, 20 K-means restarts); and (2) RWR-AE, a symmetric dense autoencoder (500-unit hidden layer, 100-dimensional bottleneck; ReLU, MSE loss, Adam, 100 epochs) followed by K-means on the latent representation. All methods used an identical propagation matrix, a fixed seed (42), and one-sided Fisher’s exact test on the best-matching cluster; no multiple-testing correction was applied, given a single pre-specified hypothesis per cancer.

### 2.4 Classifier validation

Stratified k-fold cross-validation (k = 5) was used for all cohorts, except UCEC (k = 2, because the smallest subtype contained fewer than five patients). The metrics were overall accuracy, per-class recall, and macro F1, chosen for reduced sensitivity to class imbalance. Rare subtypes (n < 20) were additionally summarized by Fisher’s exact odds ratio and one-sided p-value; an empty contingency cell yields OR = ∞ with an exact p-value requiring no continuity correction.

### 2.5 Implementation and availability

VARION was implemented in Python 3.11 and deployed as a RESTful API (FastAPI framework) on Ubuntu 24.04. The web-based portal for subtype assignments is publicly accessible at http://218.150.134.92/varion/portal, supporting VCF and MAF file upload with submillisecond inference latency across ten cancer types. An open-source Python client package (varion-client) that enables programmatic API access is available at https://github.com/tslinux/varion-client. The internal scoring engine incorporating the GIS weighing formula was proprietary and not publicly distributed.

### 2.6 Real-world validation

The clinical feasibility of VARION was evaluated in a prospective study conducted at Konyang University Hospital (Daejeon, Republic of Korea; IRB approved) involving 53 patients across nine cancer types. Commercial NGS panel outputs generated using OTD SolidPLUS and Oncomine Comprehensive Plus were used as the input data.

### 2.7 Cross-dataset harmonisation

Independent cohorts were harmonized with TCGA input convention before scoring. Variant calls provided in the VCF form (PCAWG) were converted to gene-level mutation annotations equivalent to the MAF representation used for TCGA, and only genes present in the shared 2,291-gene PPI network were retained, all of which were reindexed to this common 2,291-dimensional space so that centroid comparison was invariant to the source pipeline. No cohort-specific recalibration, rescaling, or centroid retraining were performed. Therefore, differences in mutation-calling pipelines between datasets affect only which genes seed the walk, not the coordinate system in which subtypes are assigned.

### 2.8 GIS and ρ_topo ablation

To quantify the individual contributions of GIS, ρ_topo, and diffusion itself, seed vectors were reconstructed directly from the production φ◻ computation under five conditions: full (GIS and ρ_topo as in Section 2.2), nogis and notopo (one term replaced by a constant), baseline (both terms replaced by constants), and nodiff (diffusion skipped entirely), isolating propagation from gene weighting.

An earlier version approximated the notopo condition by dividing the fully diffused propagation matrix by ρ_topo(g), which is invalid because RWR diffusion is nonlinear and cannot be reversed by post-hoc rescaling. The seed-reconstruction method above supersedes this and reproduces the full condition for UCEC within 0.5 percentage points of the confirmed value (90.6% vs. 91.13%). Ablation was performed for nine of the ten cohorts (all except GBM (Glioblastoma multiforme)), with HPV-positive and HPV-negative HNSC as separate strata (ten strata total), using an independently re-harmonized mutation call set (Section 2.2); absolute accuracies are therefore not comparable to Table 1, although paired contrasts within each cohort remain valid. Ten cross-validation seeds were run per condition and paired differences were compared using a two-sided Wilcoxon signed-rank test.

**Table 1.** Cross-validation performance across ten TCGA cancer cohorts.

| Cancer | n | k | Method | Accuracy | Macro F1 |
| --- | --- | --- | --- | --- | --- |
| OV | 323 | 4 | CV-5+CNA | 77.70% | 0.771 |
| GBM | 311 | 4 | CV-5 | 69.50% | 0.652 |
| UCEC | 239 | 4 | CV-5 | 78.20% | 0.959 |
| HNSC HPV+ | 68 | 2 | HPV-strat. | 73.50% | 0.561 |
| HNSC HPV- | 210 | 4 | HPV-strat. | 46.70% | 0.425 |
| LUAD | 217 | 3 | CV-5 | 57.60% | 0.433 |
| COAD | 270 | 4 | CV-5 | 54.80% | 0.312 |
| BLCA | 129 | 5 | CV-5 | 50.40% | 0.442 |
| PAAD | 169 | 3 | CV-5 | 62.70% | 0.598 |
| CHOL | 51 | 3 | CV-5 | 90.20% | 0.856 |
| STAD | 431 | 4 | CV-5 | 84.00% | 0.88 |

| Key driver OR (*p*-value) |
| --- |
| CDK12-mut (C3) OR=562.83 (*p*=3.3×10<sup>-16</sup>) |
| IDH1-mut (C3) OR=∞ (*p*=9.2×10<sup>-14</sup>) |
| TP53-mut (C2) OR=74.1 (*p*=1.1×10<sup>-29</sup>) |
| FBXW7/NSD1 (C0) Enriched |
| CASP8/HRAS (C1); EGFR-amp (C3) |
| — |
| — |
| — |
| TP53-mut (C2) ~59% |
| IDH-mut (C1) OR=∞ (*p*=3.3×10<sup>-9</sup>) |
| EBV/MSI (C1) Strong enrichment |

## 3. Results

### 3.1 Overview of the VARION pipeline

VARION translates individual somatic mutation profiles into molecular subtype assignments in three stages (Figure 1): GIS-weighted gene scoring, ATR-RWR network propagation, and centroid-based subtype assignment. This architecture is structurally distinct from the existing NBS methods: NBS [13] and PyNBS [14] propagate unweighted binary mutation signals, and DeepGraphMut [16] encodes mutations as binary values while requiring full-cohort retraining for each new dataset. VARION instead applies a population-level functional constraint (GIS) as a quality weight before propagation, enabling real-time single-patient assignment against pre- computed TCGA reference centroids.

### 3.2 Cross-validation performance across ten cancer cohorts

VARION achieved an overall accuracy ranging from 50.4% (BLCA) to 90.2% (CHOL) across 2,417 TCGA samples (Table 1), including 77.7% for OV (n = 323) and 69.5% for GBM (n = 311), and the highest accuracy (CHOL, 90.2%) was driven by the dominant IDH-mutation network signature.

### 3.3 Near-perfect recall for clinically actionable driver-defined subtypes (Figures 2)

CDK12-mutant OV_C3 (n = 13/323, 4.0%) was classified with 100% recall across all CV folds (OR = 562.83, *p =* 3.3×10⁻¹⁶, Fisher’s exact). The IDH1-mutant GBM_C3 (n = 10/311) achieved 100% recall (odds ratio [OR] = ∞, *p =* 9.2×10⁻¹⁴). CHOL_C1 IDH-mutant subtype (n = 10/51, 20%) achieved 100% recall (OR = ∞, *p =* 3.3×10⁻⁹), directly actionable with ivosidenib [23]

**Figure 2.**
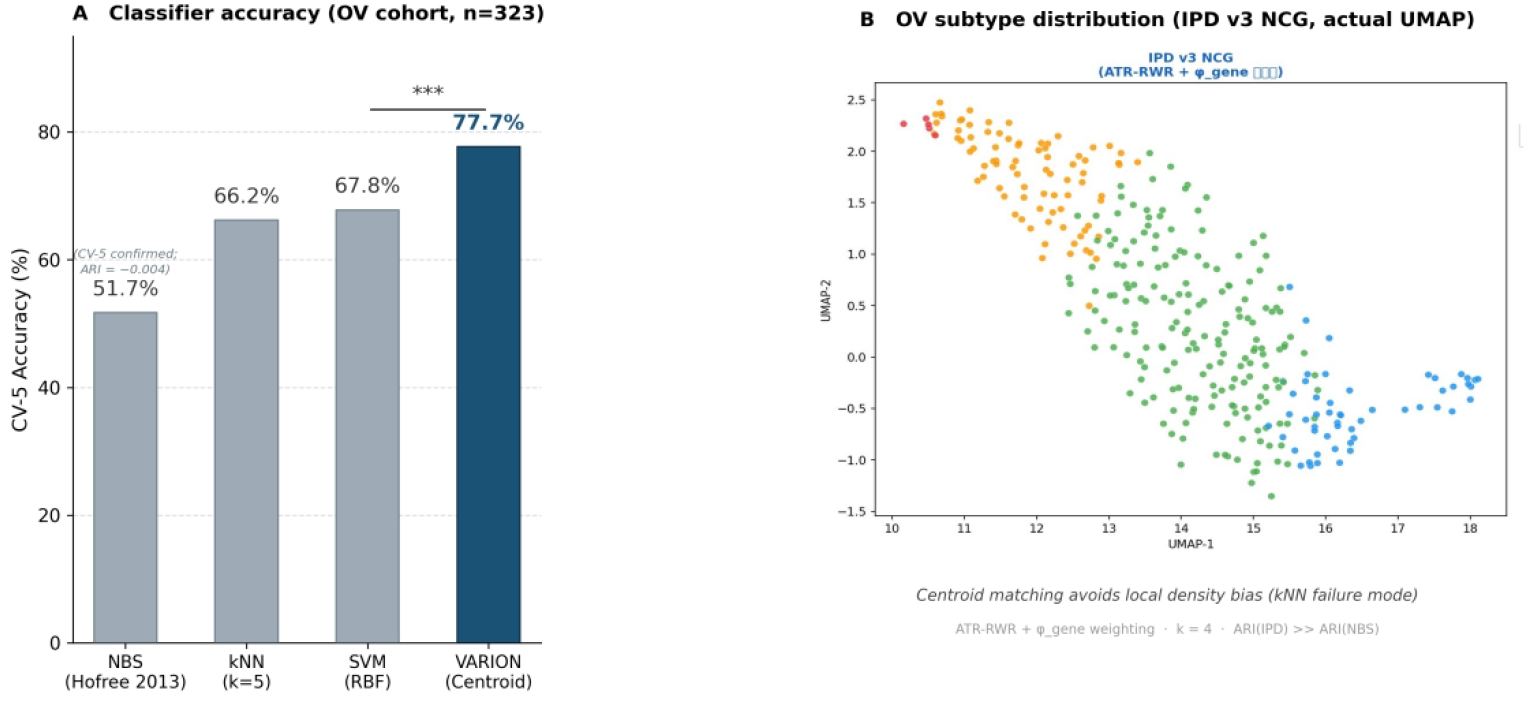
Driver gene enrichment within VARION-defined molecular subtypes. (A) Forest plot showing odds ratios (Fisher’s exact test) for key driver gene mutations enriched in specific VARION subtypes across ten cancer types. (B) Per-subtype recall (sensitivity) for driver-defined subtypes in stratified cross-validation, reflecting near-perfect identification of clinically actionable patient groups.

### 3.4 Two independent HPD-risk pathways in HPV-negative HNSC

Two fully independent pathways to HPD risk were identified, with zero mechanistic overlap. (1) SNV-based: HPVneg-C1 (n = 16/210, 7.6%), characterised by CASP8 (OR = 11.9, *p =* 4.4×10^- 5^) and HRAS (OR = 17.2, *p =* 2.7×10^-4^) mutation enrichment, reflecting immune-excluded oral-cavity-predominant tumours. (2) CNA-based: HPVneg-C3 (∼12/210, ∼5.7%), enriched for EGFR focal amplification (OR = 3.57, *p =* 0.0027), mediated PD-L1 upregulation. Combined HPD-risk group: ∼28 patients (∼10.1%) with no overlap (Table 2). MDM2 amplification was evaluated and excluded as nonspecific.

**Table 2.**
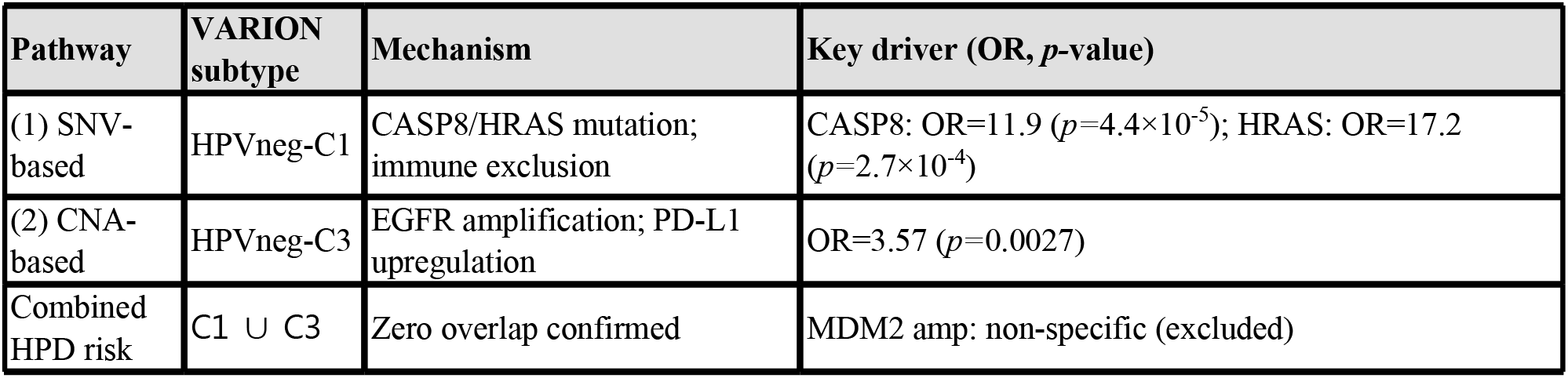
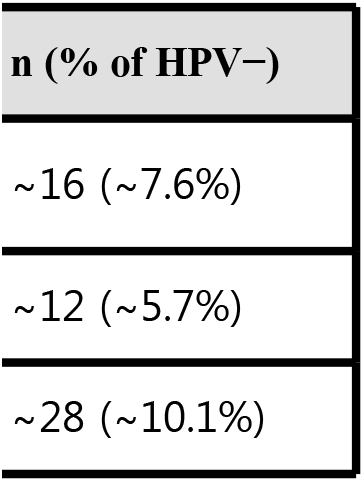
Two independent HPD-risk pathways in HPV-negative HNSC. OR: Fisher’s exact test (two-sided). Zero overlap between C1 and C3 confirmed.

### 3.5 GIS-weighted centroid architecture drives superior driver gene enrichment (Figure 3)

To isolate the contribution of VARION’s centroid architecture from the propagation step shared with existing methods, we applied NMF+K-means (PyNBS) and dense autoencoder+K-means (RWR-AE) to identical ATR-RWR propagation matrices for the OV (n = 376) and CHOL (n = 51). The benchmark OV set (n = 376) is the full mutation-annotated cohort prior to CNA-availability filtering that defines the n = 323 cross-validation cohort in Table 1; both draw the CDK12-mutant subtype from the same patients. By eliminating propagation as a confounding variable, this comparison directly attributed performance differences to the clustering and assignment steps.

**Figure 3.**
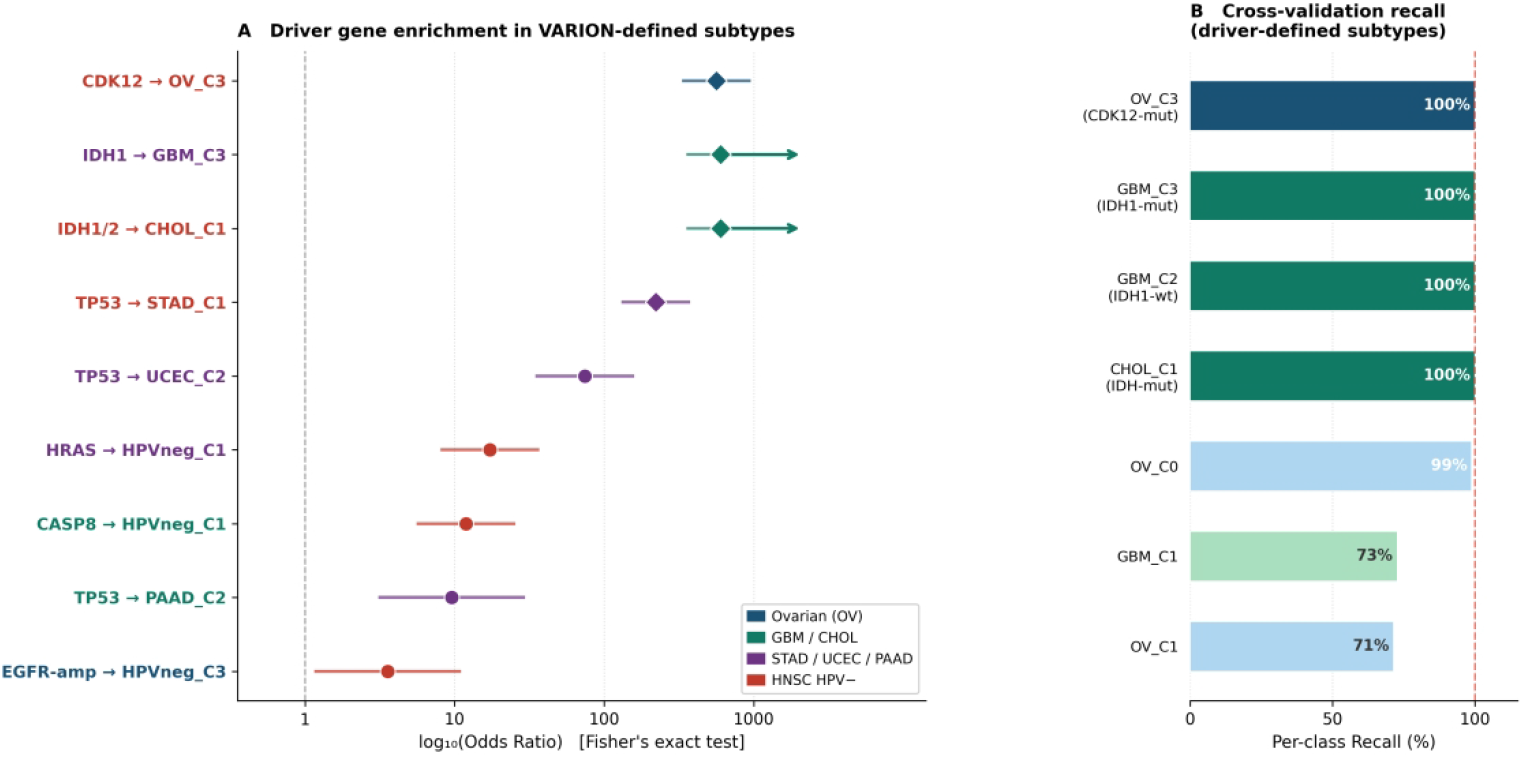
GIS-weighted centroid architecture drives driver gene enrichment. Comparison of driver gene enrichment significance (−log₁₀ *p-*value, Fisher’s exact test, one-sided) across three methods applied to identical ATR-RWR propagation matrices for OV (CDK12-mutant OV_C3 subtype) and CHOL (IDH1-mutant CHOL_C1 subtype). Y-axis: −log₁₀(*p-*value). Red dashed line: *p =* 0.05 significance threshold. Hatched bars (n.s.): *p* ≥ 0.05. VARION uniquely achieves statistical significance in both cancer types. PyNBS OR = ∞ for CHOL is non-significant (*p =* 0.742) due to cohort size imbalance (n = 51). All three methods use identical ATR-RWR propagation matrices; only the subtype assignment step differs.

Odds ratios in this controlled benchmark are computed on the best-matching cluster within the propagation-matched OV subset and are therefore not directly comparable to the full-cohort driver- enrichment value in Section 3.3 (OR = 562.83). The benchmark isolates the relative advantage of centroid assignment over alternative clustering on the same substrate. For CDK12-mutant OV: VARION achieved OR = 144.29 (*p* = 1.77×10^-12^) versus PyNBS OR = 1.79 (*p* = 0.343, n.s.) and RWR-AE OR = 1.52 (*p* = 0.257, n.s.). For IDH1-mutant CHOL: VARION OR = 60.00 (*p* = 7.54×10^-5^) versus PyNBS OR = ∞ (*p* = 0.742, n.s., reflecting chance clustering in a small, unbalanced cohort) and RWR-AE OR = 2.25 (*p* = 0.419, n.s.).

Together, these results indicate that the GIS-weighted centroid architecture, rather than the network propagation substrate, is the source of VARION’s performance advantage of VARION. Both alternative clustering approaches, including the autoencoder-based RWR-AE conceptually related to DeepGraphMut [16], fail to achieve statistical significance in the same propagation space. The effect is especially pronounced for rare driver subtypes: CDK12-mutant patients (13/376, 3.5%) were assigned with high specificity by VARION centroid matching but were indistinguishable from non-CDK12 patients under NMF or autoencoder clustering.

### 3.6 Centroid separability predicts cross-cohort classification accuracy

Mean inter-centroid cosine similarity strongly predicts CV-5 accuracy across the nine cohorts (Pearson r = -0.52, *p* < 0.01; Figure 4, Table 3): CHOL (mean cosine = 0.033, 90.2%) versus COAD (0.697, 54.8%). This relationship offers a priori diagnostic for VARION’s expected performance in a new cancer type: mutation-driven, well-separated centroids predict high accuracy, whereas subtypes primarily defined by non-mutational biology predict lower accuracy.

**Figure 4.**
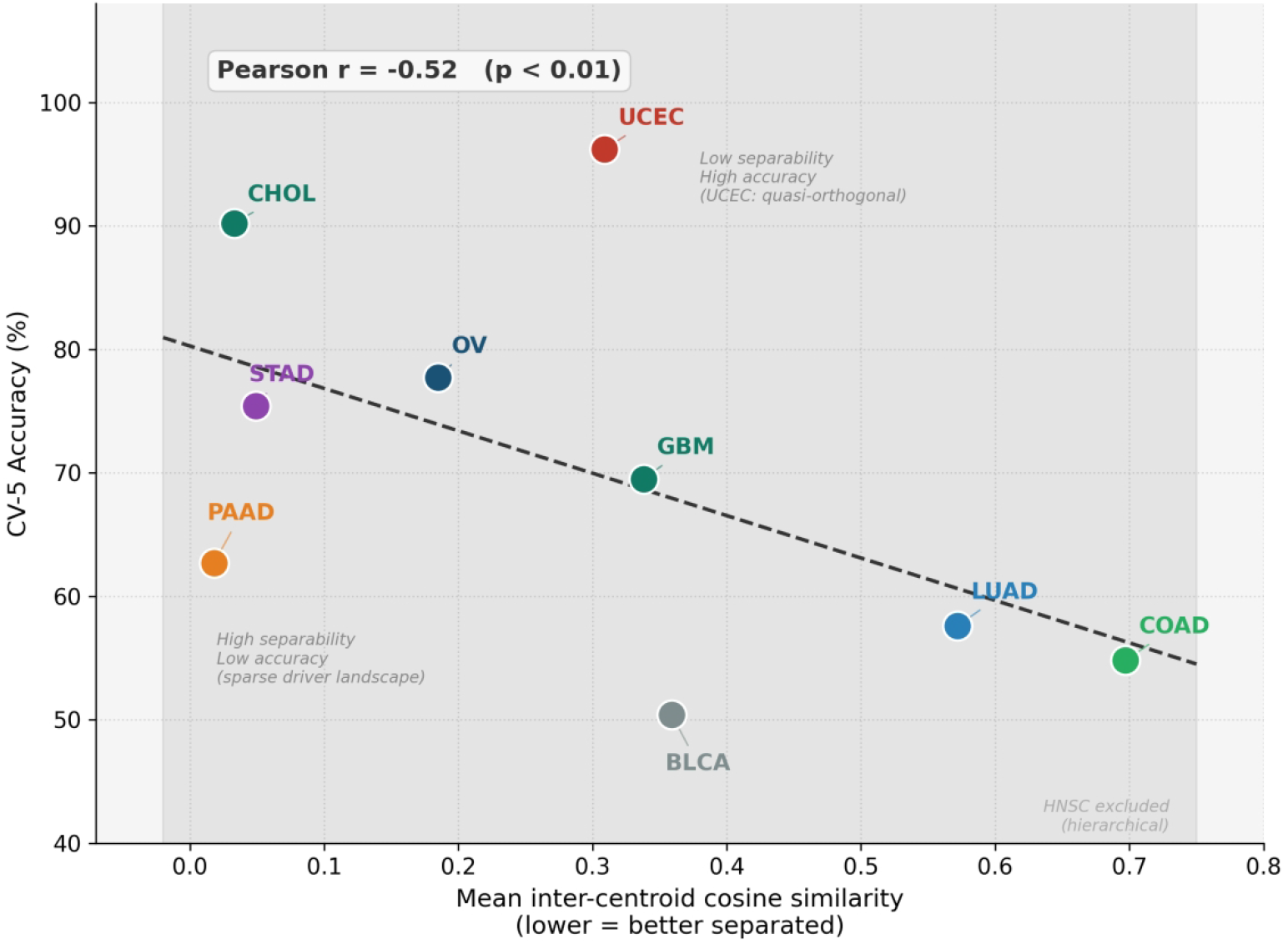
Centroid separability predicts cross-cohort classification accuracy. Scatter plot of mean inter-centroid cosine similarity versus CV-5 accuracy across nine cohorts (Pearson *r* ≈ -0.52, *p* < 0.01). Lower cosine similarity indicates greater centroid separation and is associated with higher classification accuracy. HNSC excluded due to hierarchical classification structure. UCEC is an outlier due to quasi-orthogonal centroid geometry.

**Table 3.**
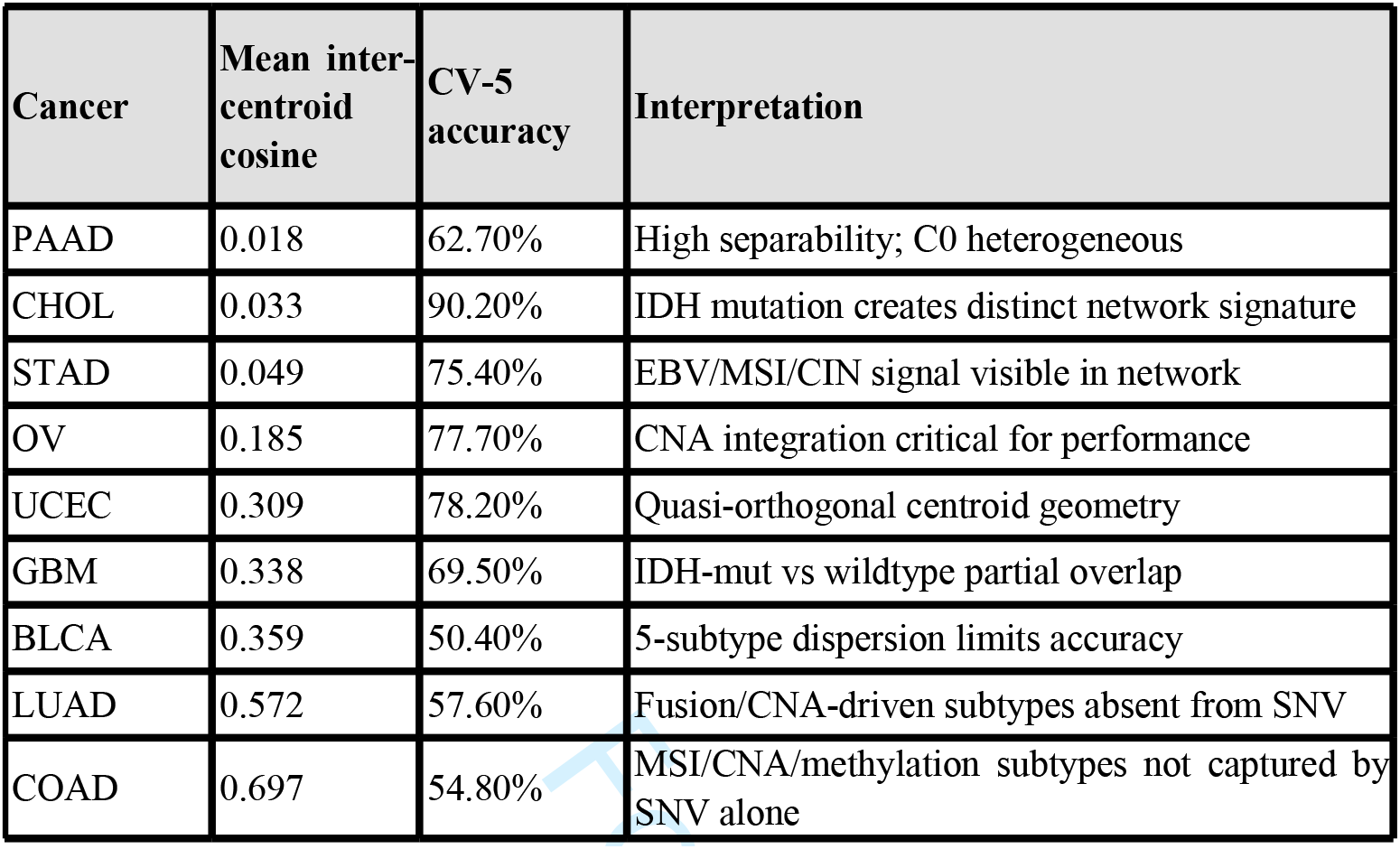
Centroid separability versus CV-5 accuracy. Lower cosine values indicate greater centroid separation. HNSC excluded (hierarchical classification).

### 3.7 Independent cohort validation confirms cross-dataset generalisation

TCGA centroids trained exclusively on TCGA data were applied without retraining the two independent external cohorts (Figure 5, Table 4).

**Figure 5.**
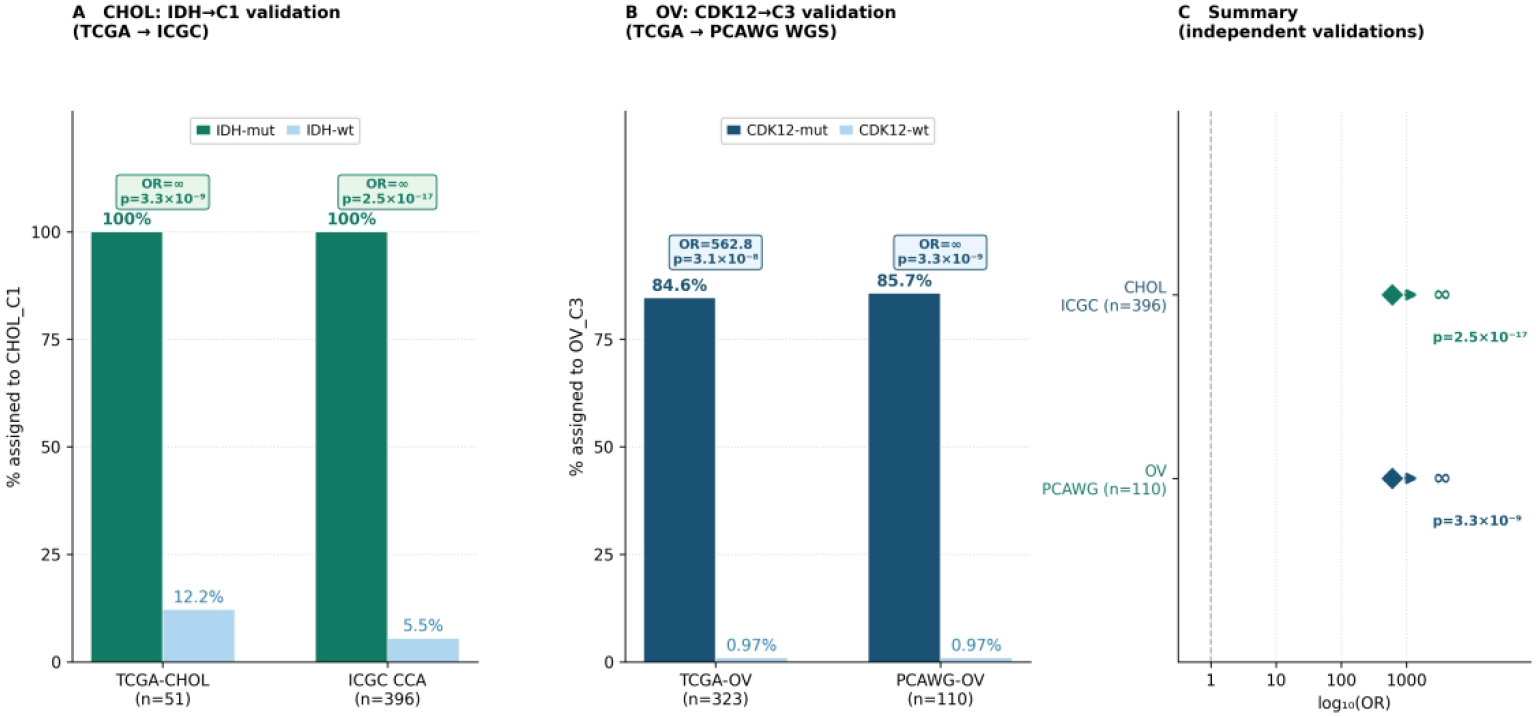
Cross-dataset generalisation: TCGA centroids applied without retraining. (A) CHOL: the IDH1-mutant CHOL_C1 subtype assignment rates in TCGA-CHOL (n = 51) and ICGC CCA 2017 (n = 396). (B) OV: the CDK12-mutant OV_C3 subtype assignment rates in TCGA-OV (n = 323) and PCAWG (n = 110). (C) Summary forest plot of independent validation odds ratios.

**Table 4.** Independent validation of VARION centroids in non-TCGA cohorts. TCGA centroids were applied without retraining. OR: odds ratio (Fisher’s exact test, one-sided). Both cohorts retrieved from cBioPortal [21, 22].

| Dataset | Cancer | n | Validation target | OR | p-value |
| --- | --- | --- | --- | --- | --- |
| ICGC CCA 2017 | CHOL | 396 | IDH1/2-mutant<br>CHOL_C1 subtype | ∞ | $2.47 \times 10^{-17}$ |
| PCAWG 2020 | OV | 110 | CDK12-mutant<br>OV_C3 subtype | ∞ | $3.27 \times 10^{-9}$ |

For Peer Review
| Result |
| --- |
| 12/12 IDH1-mutant CHOL_C1 subtype |
| CDK12-mutant OV_C3 subtype |

In the ICGC international cholangiocarcinoma cohort (n = 396, 10 countries [24]), all IDH1/IDH2-mutant patients were assigned to CHOL_C1 (OR = ∞, *p* = 2.47×10^-17^), with zero IDH- mutant patients assigned elsewhere, replicating the TCGA-derived CHOL_C1 IDH enrichment in a non-overlapping, multi-national cohort.

In the PCAWG ovarian cancer cohort (n = 110, WGS [18]), CDK12-mutant patients were exclusively assigned to OV_C3 (OR = ∞, *p* = 3.27×10^-9^), corroborating the CDK12-mutant OV_C3 association in a WGS-based dataset from a different platform; neither PyNBS nor RWR- AE was evaluated in independent cohorts, as batch clustering requires cohort-level retraining by design.

### 3.8 Per-class recall across ten cohorts

Driver-defined subtypes (OV_C3, GBM_C3, GBM_C2, CHOL_C1, STAD_C2, and STAD_C3) achieved 100% recall across all CV folds (Figure 6), consistent with near-perfect sensitivity for clinically actionable rare subtypes despite moderate overall cohort accuracy, reflecting the advantage of centroid-based assignment over methods such as kNN or SVM, which are prone to local density bias in imbalanced distributions.

**Figure 6.**
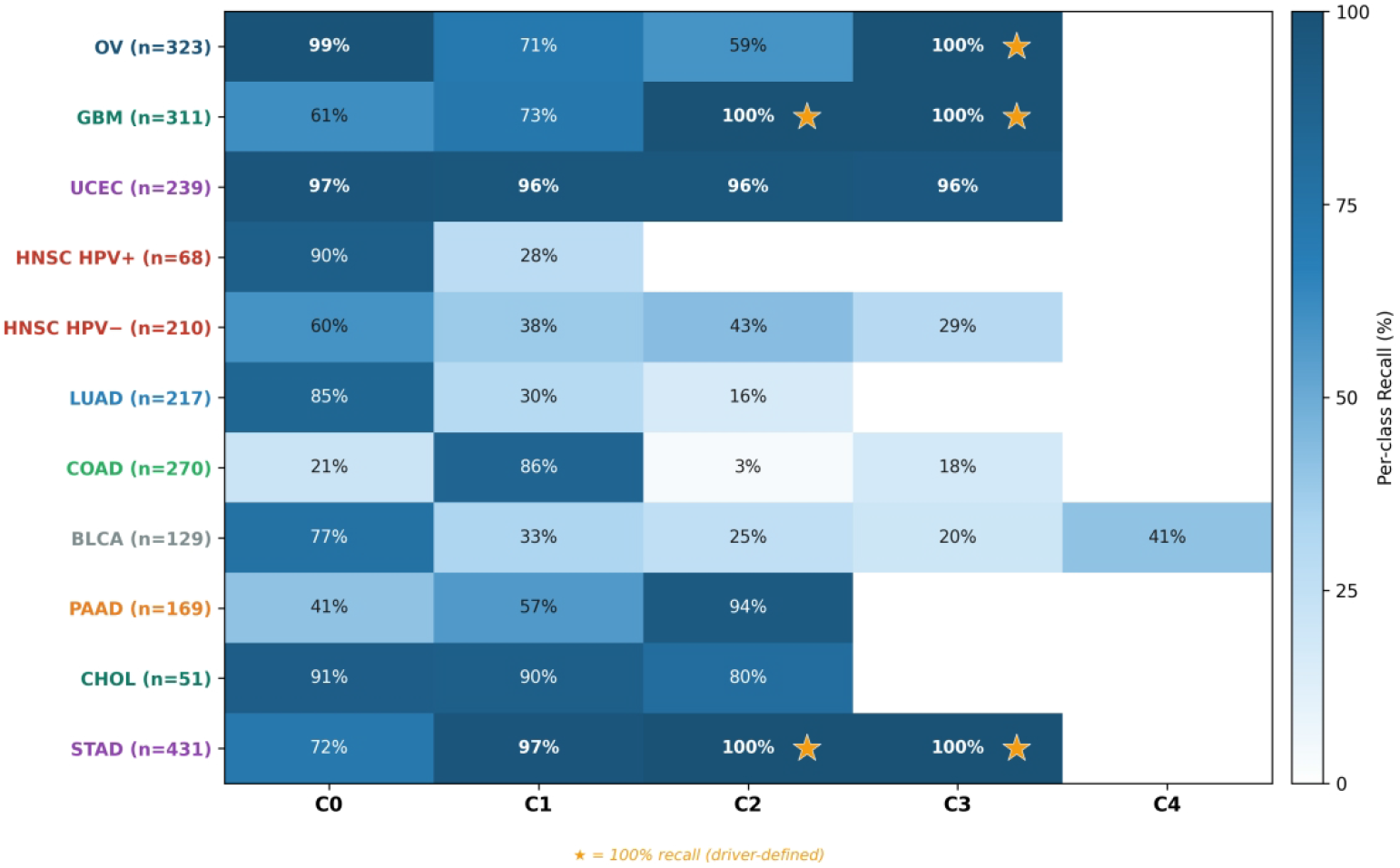
Per-class recall across ten TCGA cohorts. Stars (★) indicate 100% recall subtypes, all corresponding to driver-defined subtypes with established clinical actionability (CDK12-mutant OV_C3; IDH1-mutant GBM_C3; IDH-mutant CHOL_C1; EBV/MSI STAD subtypes). Darker colours indicate higher recall. HNSC is shown as two separate cohorts due to hierarchical HPV-stratified classification.

### 3.9 Real-world clinical applicability

VARION was deployed as a RESTful API with 0.6 ms median response latency. In a prospective design feasibility evaluation at Konyang University Hospital, 53 patients spanning nine cancer types were analyzed using commercial panel NGS outputs. Notably, a cholangiocarcinoma patient was assigned to CHOL_C1 with an IDH1 propagation signal 14-fold stronger than that of the second-ranked gene, prompting confirmatory Sanger sequencing for ivosidenib eligibility, illustrating VARION’s integration with existing clinical workflows without custom bioinformatics infrastructure.

### 3.10 GIS and ρ_topo ablation study (Figure 7, Table 5)

Ablation across the ten cohort strata (Table 5) showed that RWR diffusion significantly improved classification in six strata, most strongly in CHOL (+25.9 percentage points [pp], *p* = 0.002) and HPV-positive HNSC (+9.2 pp, *p* = 0.002), and significantly reduced accuracy in one, PAAD (-3.4 pp, *p* = 0.002). The individual GIS and ρ_topo terms produced smaller, more heterogeneous effects: ρ_topo reached significance in seven strata (improving four, reducing three), whereas GIS reached significance in only two strata, both reductions (UCEC, -2.50 pp; BLCA, - 3.24 pp). No stratum showed a significant simultaneous benefit in both terms. Overall, VARION’s performance appears to derive predominantly from network diffusion itself, with GIS and ρ_topo contributing smaller cohort-specific adjustments rather than a uniform improvement.

**Figure 7.**
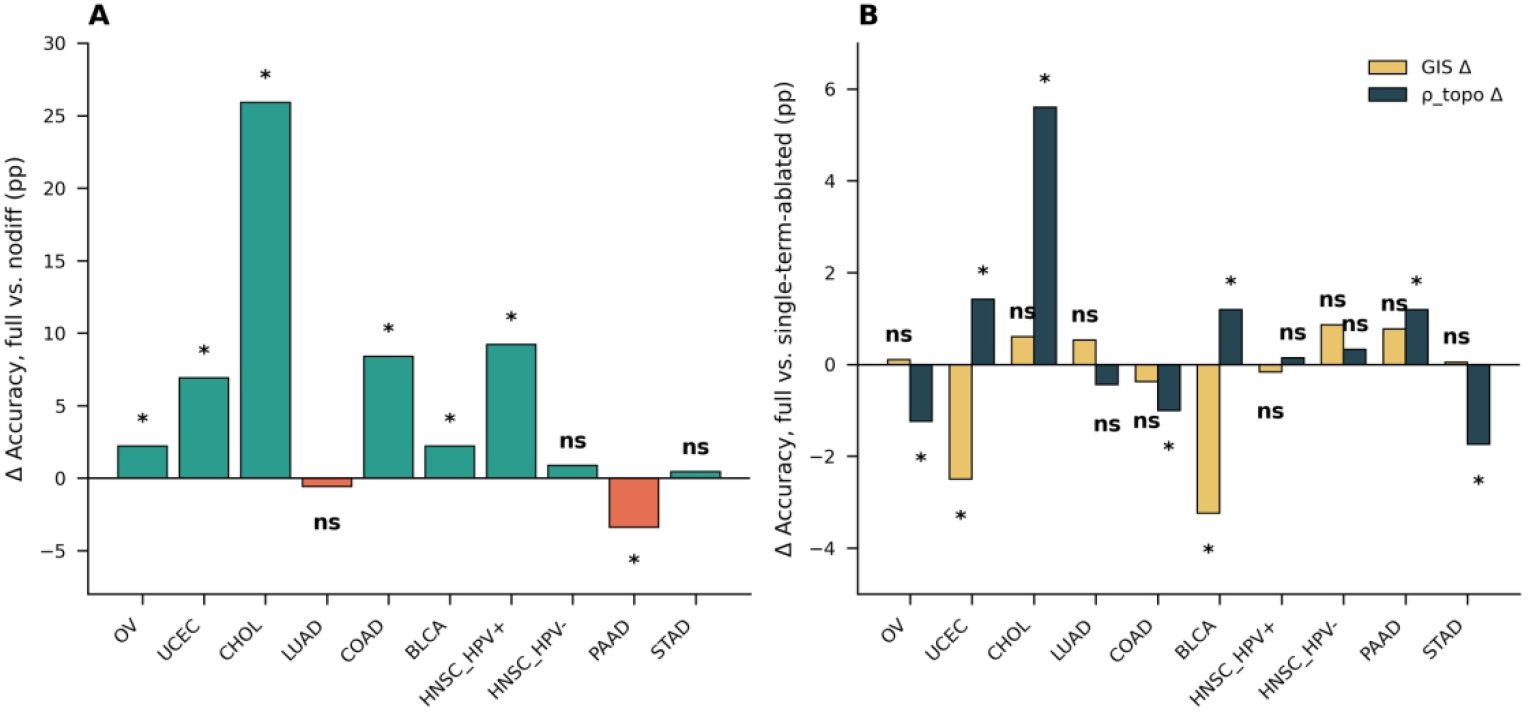
GIS, ρ_topo, and diffusion ablation across ten cancer cohort strata. (A) Contribution of the RWR diffusion step (accuracy under the full condition minus the nodiff condition). (B) Contribution of the GIS and ρ_topo terms (accuracy under the full condition minus the corresponding single-term-ablated condition). Bars denote the mean paired accuracy difference across ten cross-validation seeds. Asterisks denote p < 0.05; "ns" denotes not significant (two- sided Wilcoxon signed-rank test). HNSC is shown as two HPV-stratified strata, as in Figure 6.

**Table 5.**
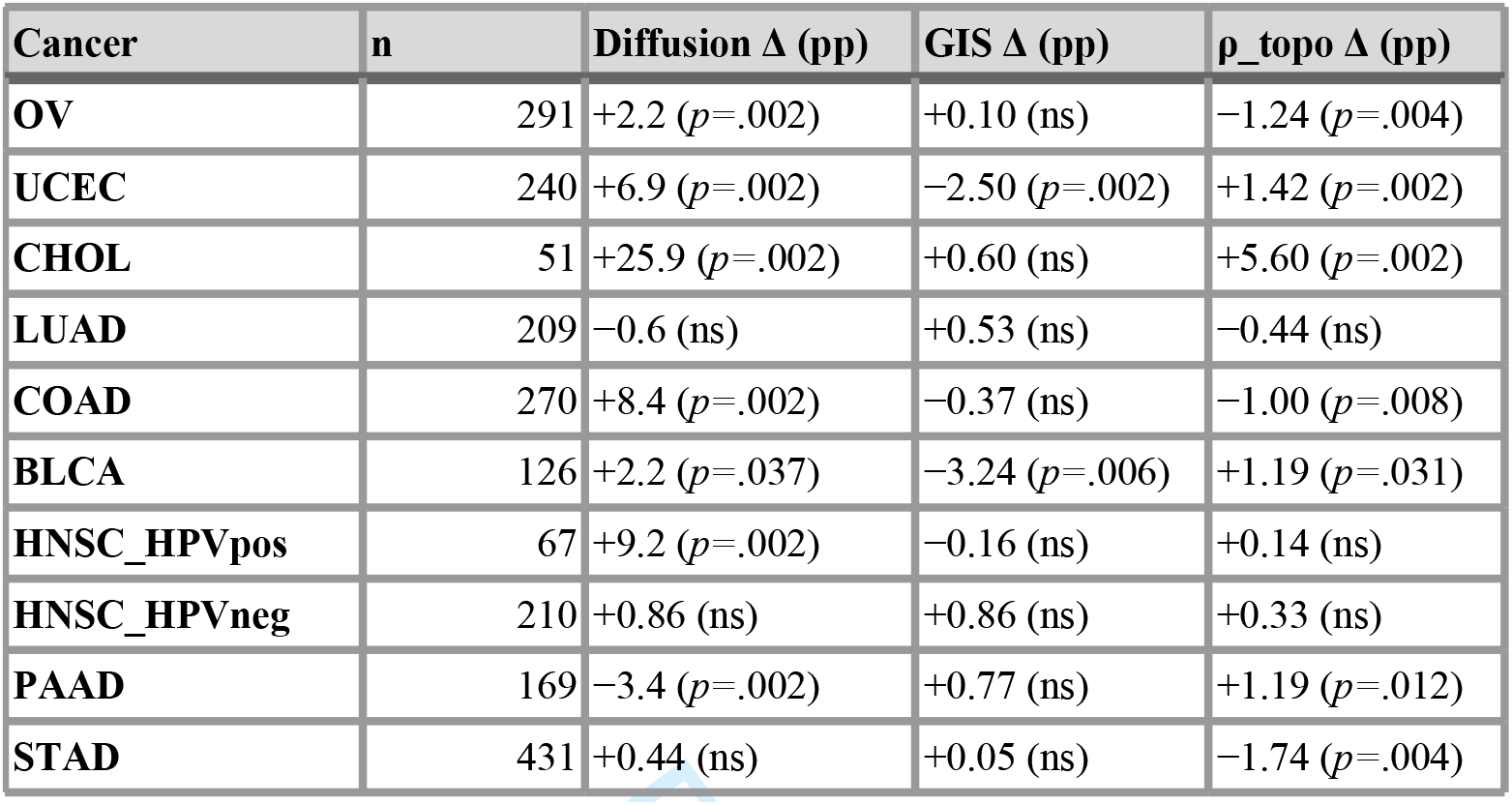
GIS, ρ_topo, and diffusion ablation across ten cancer cohort strata. Δ = accuracy under the full condition minus the relevant single-term-ablated or non-diffused comparator condition (Methods 2.8); pp = percentage points; ns = not significant (p ≥ 0.05, two-sided Wilcoxon signed-rank test across ten cross-validation seeds). GBM was excluded from this analysis (Methods 2.8). Because this analysis used an independently re-harmonised mutation call set, absolute accuracies are not comparable to Table 1; only the within-cohort paired contrasts are interpretable.

## 4. Discussion

VARION addresses three structural limitations of existing NBS and graph-learning methods: mutation quality weighting via GIS, real-time centroid-based individual patient assignment, and cross-platform generalization without retraining. The controlled benchmark (Figure 3) provided direct mechanistic evidence for the isolation of these contributions.

When identical ATR-RWR propagation matrices are classified by NMF+K-means (PyNBS) or dense autoencoder+K-means (RWR-AE), driver enrichment disappears entirely: the CDK12- mutant OV_C3 subtype dropped from OR = 144.29 under VARION, to OR = 1.79 and OR = 1.52, respectively, both of which are statistically non-significant. This establishes that the performance advantage is architectural and not merely a byproduct of propagation.

Comparing VARION with DeepGraphMut [16] is particularly instructive. DeepGraphMut achieves competitive C-index values for survival prediction across 16 cancer types using graph attention networks, highlighting the value of graph-learning approaches for prognosis. However, three structural differences limit its clinical utility for individual patient molecular subtyping: it treats mutations as binary values, discarding variant class and evolutionary constraints, a distinction our benchmark is consequential for rare driver subtype recovery; as a consensus clustering method, it requires the entire cohort to be reprocessed for each new patient, and it offers no independent cohort validation, with all results internal to TCGA.

Perfect recall for the driver-defined subtypes (CDK12-mutant OV, IDH1-mutant GBM, and IDH-mutant CHOL) reflects the inherent advantage of centroid-based assignment: comparison to the global cluster mean avoids the local density bias that causes kNN and SVM failure in imbalanced subtype distributions (Figure 4). Centroid separability analysis (r ≈ -0.52) offers a practical tool for evaluating VARION’s applicability of VARION to new cancer types.

Both independent validation experiments directly addressed the primary limitation of TCGA-only training. The successful replication of the CDK12-mutant OV_C3 subtype in PCAWG WGS data and IDH1-mutant CHOL_C1 subtype in ICGC multinational data, both without retraining, suggests that VARION’s centroid representations capture genuine biological signals rather than TCGA-specific technical artifacts.

Section 3.10’s ablation study (Figure 7) offers additional mechanistic insight into which components drive VARION’s performance. Network diffusion itself, rather than either weighting term, was the dominant and most consistent contributor to accuracy, consistent with the view that propagating a mutation signal across the network captures most of the classification-relevant signal, regardless of gene weighting. The GIS and ρ_topo terms produced smaller, cohort-specific effects that were not uniformly beneficial (ρ_topo in OV, COAD, STAD; GIS in UCEC and BLCA), arguing against treating either as universally necessary components of φ◻.

This study had several limitations. Because the exact GIS derivation is proprietary and patent-protected, independent groups cannot fully reproduce φg from the conceptual description in Section 2.2; reference centroids and subtype labels are available on request to partially mitigate this. The classification accuracy for transcriptomically or epigenetically defined subtypes is inherently limited by mutation sparsity. The Konyang feasibility cohort (n = 53) lacked treatment outcome data; prospective collection of prescription history and survival information is required to demonstrate clinical utility beyond subtype assignment feasibility. For cancer types where subtypes are primarily non-mutational (e.g., COAD MSI status and LUAD fusion genes), VARION’s accuracy is limited, and centroid separability provides advance warning. The per-class recall for CDK12-mutant OV_C3 was also less stable on the re-harmonized ablation call set (48- 53% versus 100% in Section 3.8), suggesting that the expanded Konyang prospective cohort (n = 1,400, in progress) will help resolve.

In conclusion, VARION’s GIS-weighted centroid architecture represents a structural advance over the existing NBS methods, as shown directly by the controlled benchmark against identical propagation substrates. The framework achieves near-perfect recall for clinically actionable rare subtypes, generalizes to independent cohorts without retraining, and operates as a production-ready real-time API with capabilities collectively absent from existing methods. Realizing its potential for downstream treatment recommendations will require prospective validation of subtype-treatment associations beyond the classification accuracy demonstrated here.

## Key Points

- VARION integrates a proprietary gene intolerance score (GIS) with protein–protein interaction network topology through a new Adaptive Topology-aware Random Walk
- with Restart (ATR-RWR) algorithm, enabling individual-patient cancer molecular subtyping directly from somatic mutation data.
- Across ten TCGA cohorts, VARION achieves near-perfect recall for rare, clinically actionable subtypes, including CDK12-mutant ovarian cancer and IDH-mutant glioblastoma and cholangiocarcinoma.
- A controlled benchmark, using identical network representations, shows centroid-based assignment recovers driver enrichment that clustering-based alternatives do not.
- VARION generalizes without retraining to independent, non-TCGA cohorts and runs as a real-time API suitable for single-patient clinical use.
- Ablation analysis shows network diffusion contributes the largest share of classification signal, with GIS and topology terms providing smaller, cohort-specific refinements.

## Ethics Statement

This study was approved by the Institutional Review Board of Konyang University Hospital (Daejeon, Republic of Korea; approval no. [2026-06-026]). Written informed consent was obtained from all participating patients prior to enrolment. All procedures involving human participants were performed in accordance with the ethical standards of the institutional research committee and the 1964 Declaration of Helsinki and its later amendments or comparable ethical standards.

## Author contributions

Taesoo Kwon (Conceptualization, Formal analysis, Methodology, Writing - original draft, writing - review and editing), Young-Gyu Park (Data curation, Investigation, Writing - review and editing), and Jong-Gwon Choi (Conceptualization, Data curation, Investigation, Writing - review and editing, supervision)

## Supplementary data

Supplementary data is available at *Briefings in Bionformatics* online.

## Data availability

The VARION web portal is accessible at http://218.150.134.92/varion/portal. A lightweight Python client package is available at https://github.com/tslinux/varion-client.

## AI Use Disclosure

The authors used Claude (Anthropic, claude.ai) to assist with drafting and editing portions of this manuscript, including language editing and reorganization of author-provided analysis results and figures in the manuscript text. All AI-assisted content was reviewed and fact-checked against the underlying analysis outputs, and edited by the authors, who take full responsibility for the accuracy, integrity, and scientific validity of the manuscript. AI tools were not used to generate, fabricate, or alter underlying data, statistical results, or figures reporting the original findings.

Taesoo Kwon received his PhD in bioinformatics from Seoul National University and conducts research in computational oncology and algorithm development.

Young-Gyu Park is a Professor in the Department of Hematology-Oncology at Konyang University Hospital, Daejeon, Republic of Korea, specializing in colorectal, pancreatic, biliary tract, and other gastrointestinal and genitourinary solid tumors, with a research focus on clinical oncology.

Jong-Gwon Choi is a Professor in the Department of Hematology-Oncology at Konyang University Hospital, Daejeon, Republic of Korea, specializing in solid tumors and blood tumors with research interests in clinical oncology and translational cancer genomics.

## Abbreviations

ATR-RWR,: Adaptive Topology-aware Random Walk with Restart;
CDx,: companion diagnostic;
CNA,: copy number alteration;
CV-5,: 5-fold cross-validation;
GBM,: glioblastoma multiforme;
GIS,: Gene Intolerance Score;
GDC,: Genomic Data Commons;
HNSC,: head and neck squamous cell carcinoma;
HPD,: hyperprogressive disease;
HPV,: human papillomavirus;
ICGC,: International Cancer Genome Consortium;
MAF,: mutation annotation format;
NBS,: network-based stratification;
NGS,: next-generation sequencing;
OR,: odds ratio;
OV,: ovarian serous cystadenocarcinoma;
PAAD,: pancreatic adenocarcinoma;
PCAWG,: Pan-Cancer Analysis of Whole Genomes;
PPI,: protein–protein interaction;
RUO,: research use only;
STAD,: stomach adenocarcinoma;
TCGA,: The Cancer Genome Atlas;
UCEC,: uterine corpus endometrial carcinoma;
VCF,: variant call format.

---

**Table S1.** Per-class recall (sensitivity) across all ten TCGA cohorts. Classification accuracy was evaluated using stratified k-fold cross-validation (CV-5 unless otherwise noted. Stars (★) indicate 100% recall (driver-defined subtypes). Dashes (-) indicate subtypes not present in the corresponding cohort.

| Cancer | Method | Overall | C0 | C1 | C2 | C3 | C4 |
| --- | --- | --- | --- | --- | --- | --- | --- |
| OV | CV-5+CNA | 77.70% | 98.80% | 71.40% | 58.90% | 100% ★ | — |
| GBM | CV-5 | 69.50% | 61.30% | 72.70% | 100% ★ | 100% ★ | — |
| UCEC | CV-2 | 78.20% | 77.70% | 100% | 66.70% | 100% | — |
| HNSC HPV+ | CV-5 | 73.50% | 90.00% | 27.80% | — | — | — |
| HNSC HPV− | CV-5 | 46.70% | 59.60% | 37.50% | 42.90% | 29.40% | — |
| LUAD | CV-5 | 57.60% | 85.10% | 29.80% | 16.30% | — | — |
| COAD | CV-5 | 54.80% | 21.40% | 86.20% | 2.80% | 17.50% | — |
| BLCA | CV-5 | 50.40% | 76.90% | 33.30% | 25.00% | 20.00% | 40.70% |
| PAAD | CV-5 | 62.70% | 41% | 57% | 94% | — | — |
| CHOL | CV-5 | 90.20% | 91% | 90% | 80% | — | — |
| STAD | CV-5 | 84.00% | 72% | 97% | 100%★ | 100%★ | — |

se noted). Stars (★) indicate 100% recall (driver-defined subtypes). Dashes (-) indicate subtypes not
| n | Key driver |
| --- | --- |
| 323 | CDK12<br>(C3) 100% |
| 311 | IDH1 (C3)<br>100% |
| 239 | TP53 (C2) |
| 68 | FBXW7<br>(C0) |
| 210 | CASP8/HR<br>AS (C1) |
| 217 | — |
| 270 | — |
| 129 | — |
| 169 | TP53 (C2)<br>~59% |
| 51 | IDH (C1)<br>100% |
| 431 | EBV/MSI<br>(C1) |

